# Breadth over Depth: Predictor-Agnostic Consensus for Improved Antibody–Antigen Structure Prediction

**DOI:** 10.64898/2026.11.17.767212

**Authors:** Varun Ullanat, Abhinav Gupta, Ruijiang Li, Pep Amengual-Rigo, Norbert Furtmann, Bryan Bryson, Bonnie Berger

## Abstract

Antibody–antigen structure prediction commonly relies on sampling a single co-folding predictor deeply and selecting its highest-confidence output using model-specific confidence scores. We introduce `AbPoolRank`, a predictor-agnostic structural selector that ranks mixed pools using cross-pose agreement, within-source reproducibility and interface geometry, without native confidence, predictor identities or predictor-specific parameters. On 103 cluster-disjoint, post-2021 antibody–antigen complexes from SAbDab2, three structures each from Boltz-2, Promera, Protenix-v2, ESMFold2-Fast and OpenDDE form a 15-structure pool that reaches 58.0% medium-or-better Success@1 (DockQ ≥ 0.49), exceeding confidence-based selection from every individual predictor sampled 100 times (26–52%). Existing ranking methods did not rescue deep single-predictor sampling or surpass our selector on mixed pools, with no method exceeding 51.5% on 100 structures from one co-folder and `AbPoolRank` matching the strongest single-structure methods while significantly outperforming consensus methods. `AbPoolRank` trained without one predictor lost at most 3.0 percentage points, and the frozen model accepted full MSA-conditioned ESMFold2, a sixth predictor absent from method development, without retraining or loss of accuracy. Diversification also improved the accuracy–compute frontier, with the 15-structure pool using 1.8-fold less aggregate GPU time than the most accurate single-predictor setting in our compute benchmark and 3.3-fold less than sampling that predictor 100 times. Together, these results show that shallow sampling across complementary co-folders, coupled with a confidence-free selector, provides an accurate and computationally efficient inference strategy for antibody–antigen structure prediction.

## Introduction

Deep-learning methods have transformed protein-complex structure prediction, with recent cofolding methods capable of generating assemblies directly from sequence [1–5]. Antibody– antigen complexes, nevertheless, remain challenging: near-native binding modes may be present among sampled structures yet receive low model confidence, whereas confidently ranked poses can place the antibody at an incorrect epitope [6–9]. The prevailing inference strategy is to sample one predictor repeatedly and select its highest-confidence output, assuming that greater sampling depth will eventually yield—and correctly identify—a better structure. However, confidence scores derived from predicted alignment errors are imperfect within-target discriminators and are not directly comparable across predictor families [10, 11]. As increasingly diverse co-folding systems become available, this raises a practical question that has received little study: should a fixed inference budget be spent sampling one predictor deeply or distributed across several predictors whose errors may be complementary?

The principle of consensus prediction predates modern deep learning. Meta-predictors such as Pcons and 3D-Jury ranked models from multiple fold-recognition servers using confidence normalization and inter-model structural similarity [12, 13], whereas CONSRANK exploited conserved inter-residue contacts across conventional docking solutions [14]. Modern multi-model estimation-of-model-accuracy methods extend this principle: MULTICOM_qa_ combines pairwise structural similarity with learned interface-contact probabilities, ModFOLD-dock aggregates multiple jury and interface-quality scores, and GATE represents candidate models as a pairwise-similarity graph while sub-sampling structurally overrepresented groups [15–17]. Complementary single-model approaches learn geometric, energetic or sequence-aware representations of individual interfaces [11, 18–21]. These methods demonstrate that structural agreement and interface plausibility are informative, but they do not jointly address how a fixed generation budget should be allocated across stochastic co-folding predictors, how independent cross-predictor support should be distinguished from repeated samples within one source, or whether a learned confidence-free selector can accommodate a newly introduced co-folder without retraining. Related antibody-docking workflows have improved sampling by combining unbound antibody conformations generated by several structure predictors before information-driven docking [22], but do not pool and select end-to-end co-folded complex poses under a fixed budget.

Here we formulate inference as a fixed-budget pool-selection problem and evaluate five contemporary co-folding predictors—Boltz-2 [23], Promera [5], Protenix-v2 [24], ESMFold2-Fast [25] and OpenDDE [26]—on 103 training cluster-disjoint, post-2021 antibody–antigen complexes from SAbDab2 [27]. We introduce a predictor-agnostic structural selector called `AbPoolRank` that combines cross-pose agreement, within-source reproducibility and interface geometry without using native confidence or predictor-specific parameters. Sampling three structures from each predictor and ranking the resulting 15-structure pool using `AbPoolRank` achieves 58.0% medium-or-better Success@1 (DockQ ≥0.49) [28], exceeding every individual predictor sampled 100 times while using less aggregate GPU time. The improvement is driven primarily by broader candidate coverage rather than deeper sampling, and persists under external quality-assessment rankers, with `AbPoolRank` matching or exceeding the strongest alternatives. `AbPoolRank` also transfers to predictors excluded from training and accepts a newly introduced co-folder without retraining. Together, these results show that predictor complementarity, coupled with a confidence-free structural selector, provides a more effective inference strategy for antibody–antigen co-folding than allocating the same budget to deep sampling from a single predictor.

## Results

### Shallow multi-predictor pools outperform deep sampling from individual predictors

We evaluated structure selection on 103 post-2021 SAbDab2 [27] antibody–antigen targets for which Boltz-2 [23], Promera [5], Protenix-v2 [24], ESMFold2-Fast [25] and OpenDDE [26] each contributed 100 candidate structures (“Methods” sections “Dataset and partition” and “Structure generation”). We compared shallow pools spanning all five predictors with increasingly deep single-predictor pools, using Success@1—the fraction of targets whose selected structure reached CAPRI medium-or-better quality (DockQ≥ 0.49)—as the primary metric [28]; pools were redrawn 100 times per target, and confidence intervals were obtained by paired bootstrap resampling of targets (“Methods” section “Endpoint and statistics”). `AbPoolRank` combines within-pool ranks of geometric pose support, source-balanced support and epitope agreement, together with pool size, without using named predictor identity or native confidence (Fig. 1a; “Methods” section “`AbPoolRank` design”). With only three structures per predictor, the 5 × 3 pool reached 58.0% Success@1, exceeding all five predictors even when each was sampled 100 times (Fig. 1b,d). The strongest single-predictor baselines were OpenDDE at 51.5% and Protenix-v2 at 48.5%, giving paired gains of +6.5 percentage points (95% confidence interval, +0.5 to +12.8) and +9.4 points (+1.0 to +17.7), respectively; gains over ESMFold2-Fast, Boltz-2 and Promera were +31.7, +29.8 and +25.9 points. The 5 ×2 pool already reached 55.8%, although its margins over Protenix-v2 (+7.2; −0.8 to +15.2) and OpenDDE (+4.3; 1.6 to +10.5) were not statistically resolved. CAPRI-class decomposition further showed increased recovery of both medium- and high-quality structures, with 31% of targets receiving a high-quality structure from the 5 × 3 pool compared with at most 27% for any predictor at 100 structures (Fig. 1c).

**Figure 1:**
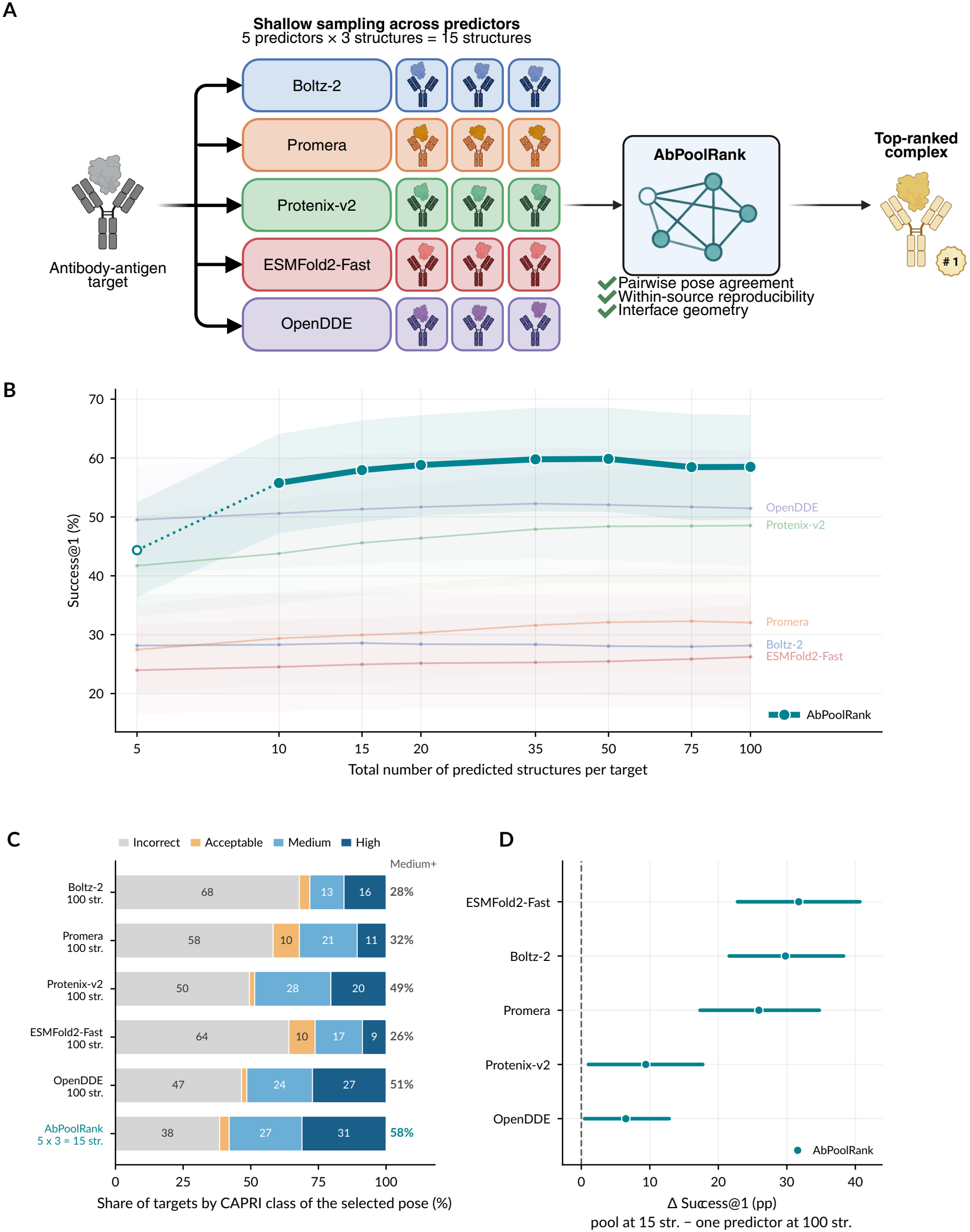
Shallow sampling across predictors outperforms deep sampling from one predictor. **a**, Method schematic. A fixed inference budget (5 predictors with 3 structures each) is distributed across co-folding predictors, and the resulting mixed pool is ranked by `AbPoolRank` using cross-pose agreement, within-source reproducibility and interface geometry. **b**, Success@1, defined as the fraction of targets whose selected structure reaches DockQ ≥ 0.49, as a function of the total number of predicted structures per target across 103 post-2021 antibody–antigen complexes. Faint colored curves show individual pre-dictors ranked by native confidence; the thick teal curve shows `AbPoolRank` applied to the five-predictor pool. Each point averages 100 redrawn pools per target; shaded bands denote marginal target-bootstrap 95% confidence intervals. The five-structure pool is shown as a hollow marker because one pose per predictor does not permit measurement of within-source reproducibility and makes source-balanced support identical to unweighted pose-similarity support; the intended operating regime therefore begins at two poses per predictor. **c**, CAPRI class of the selected structure for each predictor sampled 100 times and for `AbPoolRank` at the 5 × 3 = 15-structure operating point. Values at right margin denote medium-or-better Success@1. **d**, Paired per-target difference in Success@1 between the 5 × 3 pool and each predictor sampled 100 times. Points denote means across 103 targets and bars denote 95% paired-bootstrap confidence intervals.

Oracle and matched-control analyses indicated that this advantage primarily reflected broader candidate coverage rather than simply more samples or source labels. The oracle ceiling, which credits the best structure present in a pool (”“Methods”” section “Baselines”), was 70.5% for the 5 × 3 pool, compared with 64.1% for 100 OpenDDE structures, while structural selection from the mixed pool and native-confidence selection from OpenDDE recovered similar fractions of their respective ceilings (82% and 80%; Supplementary Fig. S1). OpenDDE supplied the highest-DockQ candidate in only 31.9% of 5 × 3 pools, indicating that most mixed pools obtained their best structure from another predictor; however, random selection from the same mixed pool reached only 30.2% Success@1. In matched controls, auxiliary predictors used only to provide agreement context did not increase the fixed primary-predictor oracle ceiling, and pseudo-source pools formed by partitioning 15 structures from a single predictor into five arbitrary groups remained below the true five-predictor pool and showed no consistent advantage over native-confidence selection (Supplementary Figs. S2 and S3; “Methods” section “Diversity controls”). Thus, the gain depended on combining distinct predictor-specific pose distributions rather than on additional candidate count or source labels alone.

Sampling depth also determined which source-aware features were available. With one structure per predictor, within-source reproducibility could not be measured and source-balanced support was identical to unweighted pose-similarity support; Success@1 was 44.3% in this setting. Increasing to two structures per predictor raised Success@1 to 55.8% and made the full source-aware representation available, while three structures per predictor increased it further to 58.0%. These results indicate that shallow sampling across distinct predictors improved structure selection relative to substantially deeper sampling from individual predictors, with three structures per source providing improvement over the strongest single-predictor baselines.

### AbPoolRank matches or exceeds existing external rankers

The advantage of the 5×3 pool could in principle reflect weak native confidence rather than improved structural selection. We therefore benchmarked published quality-assessment methods on the same candidate structures (Fig. 2; “Methods” section “External rankers”). VoroIFGNN [20], DeepRank-Ab [11] and DProQA [19] score structures independently and were applied to both deep single-predictor and mixed pools, whereas Pcons [12], 3D-Jury [13], MULTICOM pairwise similarity score (PSS) [15] and a DockQ-based jury were applied to the mixed pools. External rankers did not rescue deep sampling (Fig. 2a). On 100 OpenDDE structures, VoroIF-GNN and DeepRank-Ab reached 49.5% and 50.5% Success@1, respectively, compared with 51.5% for native confidence; on Protenix-v2 they reached 41.7% and 38.8%, compared with 48.5%. Across all 20 co-folder–ranker combinations, the best deep-sampling result remained OpenDDE with native confidence. By contrast, `AbPoolRank` reached 58.0% on only 15 mixed structures, exceeding deep OpenDDE sampling by +6.5 percentage points (95% confidence interval, +0.5 to +12.8). Its margin increased to +8.4 points over OpenDDE ranked by VoroIF-GNN and +7.5 points over DeepRank-Ab. `AbPoolRank` also exceeded the oracle ceilings of 100 Boltz-2 and ESMFold2-Fast structures by +22.0 and +20.1 points, respectively, and was level with the Promera oracle. Only the OpenDDE and Protenix-v2 oracles, at 64.1% and 62.1%, remained above it, indicating that for most co-folders the limitation of deep sampling was the candidate set itself rather than the ranking method.

**Figure 2:**
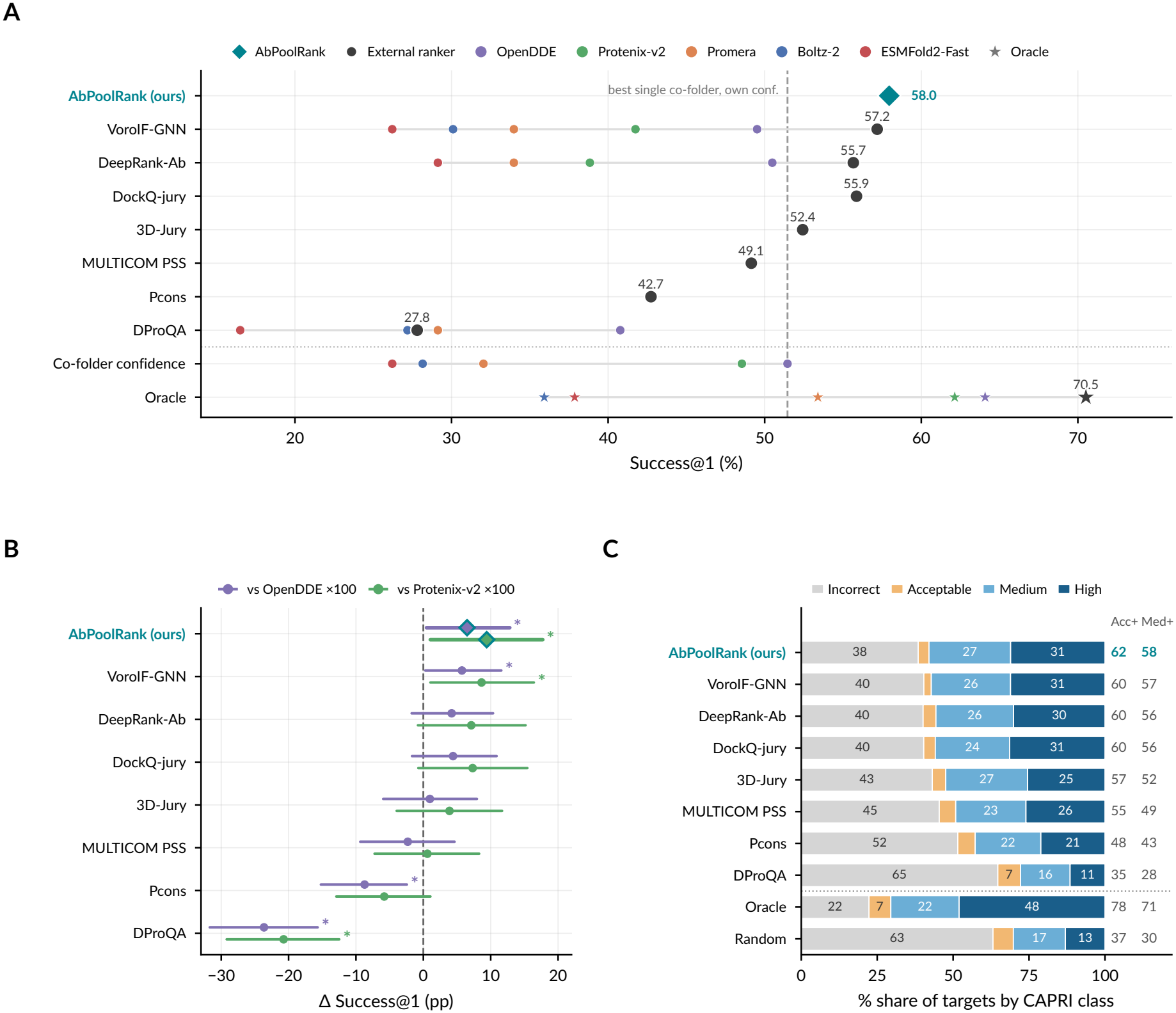
AbPoolRank matches or exceeds external rankers. All panels use the 103 test targets. Mixed-pool results use the 5 × 3 = 15-structure pools from Fig. 1, with 100 draws per target and deep-sampling results use 100 structures from one co-folder. Each method is credited with the DockQ of its top-ranked structure, with unscored structures ranked last. **a**, Success@1 for each ranking method. Large markers show performance on the 15-structure mixed pool; small colored markers show the same ranker applied to 100 structures from an individual co-folder. The co-folder confidence row uses each predictor’s native confidence. Stars denote oracle performance for the mixed pool and individual co-folders. The dashed line marks the strongest deep-sampling result, OpenDDE ranked by native confidence (51.5%). Consensus methods (DockQ-jury, 3D-Jury, MULTICOM PSS and Pcons) are defined only for pools and therefore have no single-co-folder counterpart. **b**, Paired difference in Success@1 between each method on the 15-structure mixed pool and native-confidence selection from 100 OpenDDE or Protenix-v2 structures, the two strongest single co-folders. Bars denote 95% paired target-bootstrap confidence intervals; asterisks mark intervals excluding zero. **c**, CAPRI class of structures selected from the 15-structure mixed pools: incorrect (DockQ *<* 0.23), acceptable (0.23– 0.49), medium (0.49–0.80) and high (≥0.80). Values at right margin denote acceptable-or-better (Acc+) and medium-or-better (Med+) fractions. Oracle and random selection from the same pools are shown below the dotted line.

`AbPoolRank` also matched or exceeded external rankers on the mixed pools (Fig. 2b). Pooling improved performance for other rankers that tracked interface quality: VoroIF-GNN on the 15-structure mixed pool outperformed VoroIF-GNN on 100 structures from every individual co-folder by +7.7 to +31.0 points, and DeepRank-Ab showed the same pattern for four of five co-folders. DProQA was the exception; its scores were nearly uncorrelated with DockQ within targets (median Spearman *ρ* = 0.05, versus 0.37 for VoroIF-GNN and 0.36 for DeepRank-Ab), and its mixed-pool performance remained poor. Among methods applied to the same 5 × 3 pools, `AbPoolRank` achieved the highest Success@1 at 58.0%. It exceeded all four consensus methods, with gains ranging from +2.1 points over the DockQ-jury to +15.2 points over Pcons, and exceeded DProQA by +30.1 points. Its Success@1 was also numerically higher than VoroIF-GNN and DeepRank-Ab by +0.8 and +2.3 points, respectively. The CAPRI-class distribution showed the same pattern (Fig. 2c): `AbPoolRank` produced the lowest fraction of incorrect predictions (38.5%) and the highest fraction of acceptable-or-better predictions (61.5%). At high quality (DockQ ≥ 0.80), `AbPoolRank` (31.1%), VoroIF-GNN (31.1%) and the DockQ-jury (31.4%) were indistinguishable, while the pool oracle reached 48.0%, leaving substantial headroom in identifying the best available structure. Despite matching or exceeding the strongest external rankers, `AbPoolRank` was substantially cheaper to evaluate: ranking a 15-structure pool required a median of 1.2 s on one CPU core, including structure parsing and pairwise features, compared with 19.3 s for VoroIF-GNN on the same pools (“Methods” section “External rankers”). Together, these results show that `AbPoolRank` converts a shallow, heterogeneous candidate pool into predictions that match or exceed substantially more expensive external rankers, while retaining higher raw Success@1 than the strongest learned alternatives

### AbPoolRank transfers to predictors excluded from training

To test transfer across co-folders, we trained five versions of `AbPoolRank`, each excluding all structures from one predictor, froze their parameters and evaluated them on mixed pools containing all five predictors (Fig. 3a; “Methods” section “Transfer experiments”). The heldout predictor contributed neither training structures nor labels. At a 15-structure budget, every leave-one-predictor-out `AbPoolRank` model remained within 3.0 percentage points of the all-predictor reference (Fig. 3b). This reference was refitted with the same procedure and evaluated on the same pool draws as the leave-one-out models, reaching 58.2% Success@1; the released `AbPoolRank` scores 58.0% on the draws of Fig. 1. Excluding OpenDDE, Protenix-v2 or ESMFold2-Fast changed performance by only 0.2–0.3 points, and excluding Promera by 0.7 points. Only withholding Boltz-2 produced a resolved loss (−3.0 points; 95% confidence interval, −5.2 to −1.1). Thus, the structural scoring rule transferred across predictor-specific pose distributions with little dependence on labeled examples from any single co-folder.

**Figure 3:**
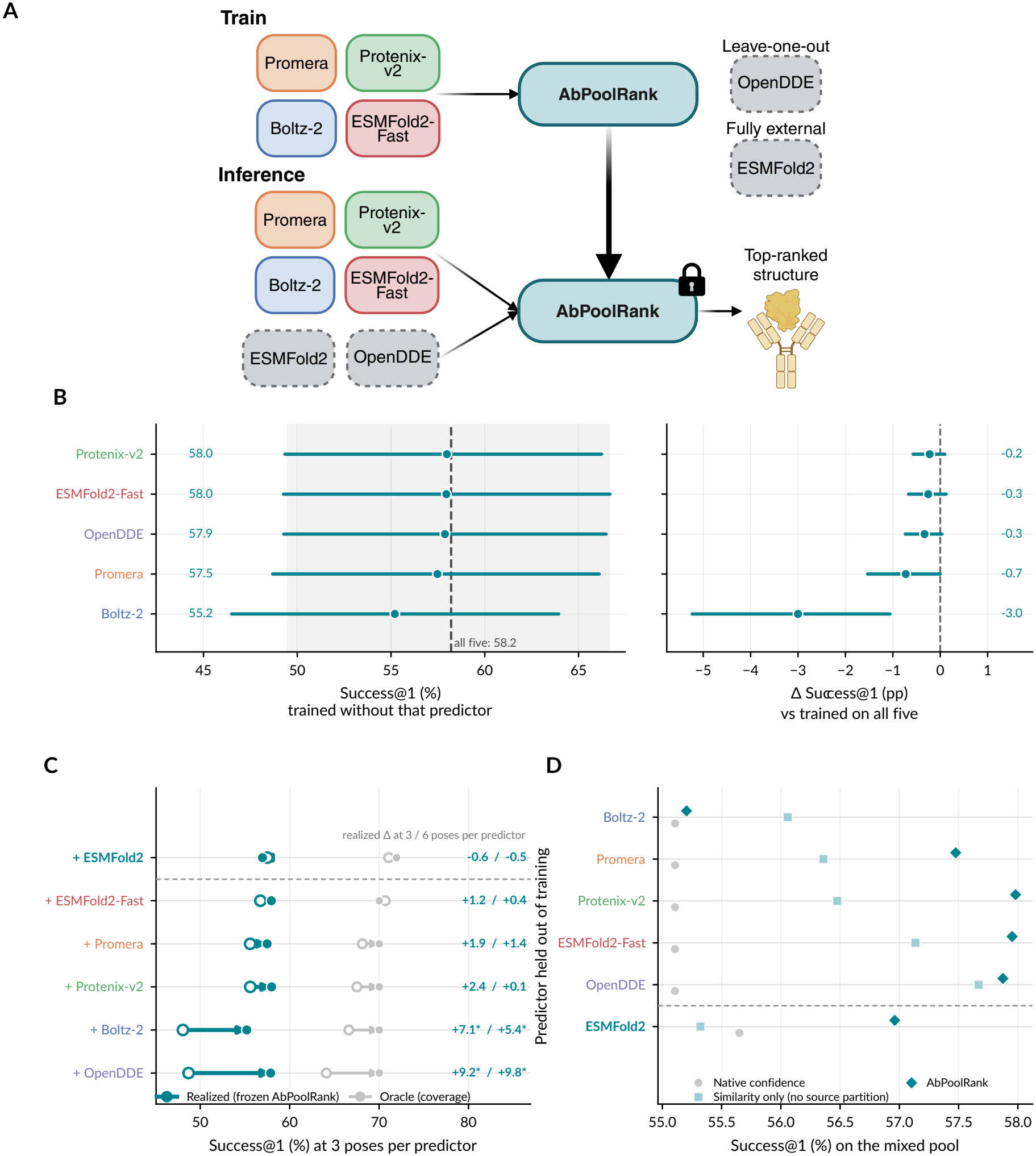
AbPoolRank transfers to predictors excluded from training. **a**, Experimental design. `AbPoolRank` is trained on a subset of predictors, frozen and evaluated on pools containing an unseen predictor. Leave-one-predictor-out models exclude one of the five development predictors from `AbPoolRank` training, whereas full MSA-conditioned ESMFold2 was absent from the development corpus. **b**, Leave-one-predictor-out transfer at a 15-structure budget. Left, absolute Success@1 for `AbPoolRank` versions trained without the indicated predictor and evaluated on pools containing it; the dashed line and grey band denote the all-five-trained reference and its marginal 95% confidence interval. Right, paired difference from the all-five-trained reference. **c**, Contribution of an unseen predictor at inference. Open circles show pools containing only predictors represented during training; filled circles show the same overall sampling after adding the unseen predictor. Teal denotes realized Success@1 under frozen `AbPoolRank` and grey denotes the pool oracle. Values at right give paired changes at three and six poses per predictor. The row below the dashed line corresponds to full MSA-conditioned ESMFold2; six-predictor pools therefore contain 18 or 36 structures, compared with 15 or 30 structures for the five-predictor leave-one-out experiments. **d**, Success@1 at three poses per predictor for native-confidence selection, a similarity-only selector that omits anonymous source membership, and `AbPoolRank`. The difference between the latter two measures the contribution of source-aware aggregation beyond global pose similarity. The row below the dashed line denotes full MSA-conditioned ESMFold2.

We next asked whether a predictor excluded from training could nevertheless improve selection when introduced at inference (Fig. 3c). At a fixed budget of 15 structures, adding the held-out predictor while redistributing samples across sources (“Methods” section “Transfer experiments”) improved Success@1 in all five cases. The largest gains came from OpenDDE (+9.2 points) and Boltz-2 (+7.1), with smaller gains from Protenix-v2 (+2.4), Promera (+1.9) and ESMFold2-Fast (+1.2). OpenDDE and Boltz-2 remained beneficial at 30 structures, whereas the other additions contributed at most 1.5 points. The value of a predictor therefore depended on its complementarity to the existing pool rather than simply on standalone accuracy: Boltz-2 had one of the lowest single-predictor oracle ceilings, yet appeared in 99% of the best three- and four-predictor subsets, and removing it from the full pool cost 7.3 points, second only to OpenDDE at 9.6 points (Supplementary Figs. S4 and S5; Supplementary Note 1; “Methods” section “Pool composition”).

We further tested a genuinely external co-folder, full MSA-conditioned ESMFold2 [25], whose structures were generated only after `AbPoolRank` and its features had been fixed (“Methods” section “Structure generation”). `AbPoolRank` accepted this predictor without modification. Full ESMFold2 was the third strongest of the six predictors by mean DockQ, and its pose distribution was not redundant with ESMFold2-Fast: their mean pairwise pose similarity (geom sim; “Methods” section “Pose-pair descriptors”) was 0.0801. Nevertheless, adding full ESMFold2 to the five-predictor pool at a fixed 18-structure budget left Success@1 unchanged within uncertainty (57.5% to 57.0%), despite a small increase in the oracle ceiling from 71.1% to 72.0%; the same pattern held at 36 structures. Thus, `AbPoolRank` transferred to a predictor absent from both training and method development without a measurable loss in accuracy, although that predictor did not improve selection beyond the existing five-model pool. Which source supplies the selected structure, and why `AbPoolRank` under-selects the external predictor relative to how often it holds the best candidate, is analyzed in Supplementary Note 2.

### Diversification improves the accuracy–compute frontier

Because co-folders differ in setup, trunk and diffusion costs, we benchmarked GPU time directly and modeled fixed setup, per-seed and per-structure costs for each predictor (Fig. 4a; “Methods” section “Compute measurement”). Setup costs varied substantially, from 0.03 min for Boltz-2 to 2.43 min for OpenDDE. Despite paying the setup cost of all five predictors, the 5 × 3 pool reached 58.0% Success@1 at 0.157 GPU-hours per target (Fig. 4b). This required 1.8-fold less aggregate GPU time than the most accurate single-predictor setting in the compute benchmark, OpenDDE with 50 structures at 52.0% Success@1, and 3.3-fold less than generating 100 OpenDDE structures. No individual predictor exceeded 52.0% at any benchmarked budget, placing the mixed pool at a more favorable accuracy–compute operating point.

**Figure 4:**
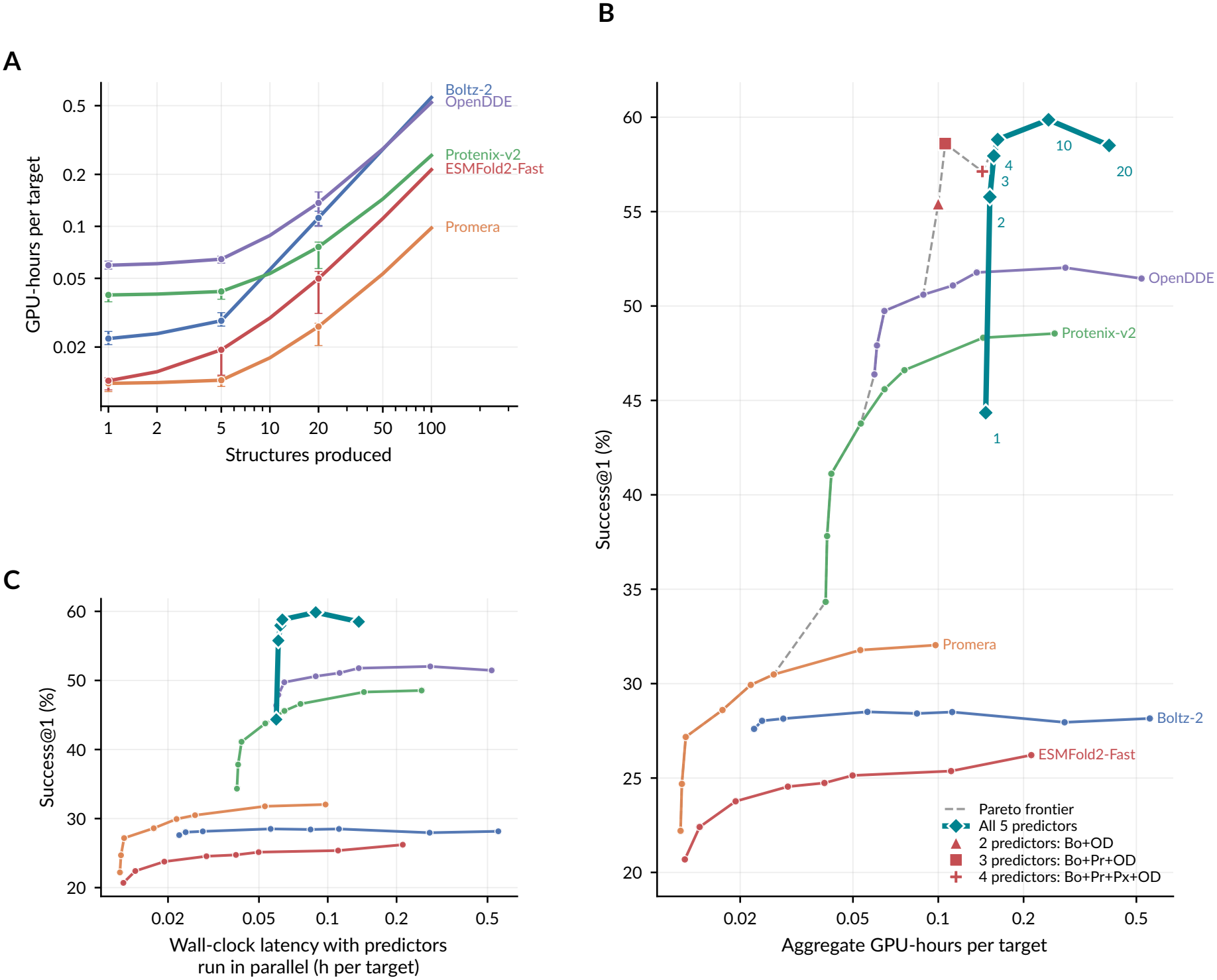
Diversification improves the accuracy–compute frontier. **a**, Measured GPU-hours per target as a function of structures generated by each predictor on an exclusive NVIDIA A100 80-GB GPU with cached MSAs. Points denote medians across eight targets near the median sequence length and error bars denote interquartile ranges. Runs were executed serially after a discarded warm-up. **b**, Success@1 against aggregate GPU-hours per target. Coloured curves show individual predictors at increasing sampling depth; teal diamonds show five-predictor pools ranked by `AbPoolRank` and are annotated by poses per predictor; red markers show smaller predictor subsets selected in the subset analysis. The dashed grey line denotes the Pareto frontier. Aggregate compute includes the fixed setup cost of every predictor in a pool. **c**, Success@1 against estimated wall-clock latency when predictors are run concurrently on separate GPUs. This represents reduced completion time through parallel execution rather than a further reduction in aggregate GPU usage.

Once the fixed cost of including multiple predictors was paid, additional depth was comparatively inexpensive. Increasing from two to three and four structures per predictor raised Success@1 from 55.8% to 58.0% and 58.8%, at 0.152, 0.157 and 0.162 GPU-hours per target, respectively; ten structures per predictor reached 59.9% at 0.245 GPU-hours, after which further sampling provided no improvement. By contrast, one structure per predictor reduced Success@1 to 44.3% while saving little compute. Every Pareto-efficient point above approximately 55% Success@1 therefore used more than one predictor. Smaller complementary subsets provided further savings when the predictor set was known in advance: at a 15-structure budget, Boltz-2, Promera and OpenDDE reached 58.6% Success@1 at 68% of the full five-predictor pool’s GPU time, placing this subset on a more favorable aggregate-compute frontier (Fig. 4b; Supplementary Note 1; “Methods” section “Pool composition”).

Diversification also reduced latency when predictors were run in parallel. With one GPU per co-folder, the 5 × 3 pool had an estimated wall-clock latency of 0.062 h, compared with 0.521 h for 100 OpenDDE structures, an approximately 8.4-fold reduction (Fig. 4c; “Methods” section “Compute measurement”). This latency gain uses additional parallel GPUs and is distinct from the reduction in aggregate GPU time. The selection model was negligible relative to structure generation, with ranking of a 15-structure pool from structure files taking about 1 s on one CPU core. Together, these results show that diversification improves the accuracy– compute frontier even after accounting for predictor-specific setup costs, while carefully chosen complementary subsets can recover similar accuracy at still lower computational cost.

## Discussion

Our results show that inference-time scaling for antibody–antigen co-folding benefits more from combining predictors than from sampling any one predictor deeply, provided that the resulting pool can be ranked effectively. Fifteen structures drawn three at a time from five predictors and ranked by `AbPoolRank` outperformed every single predictor sampled 100 times, while using 1.8-fold less aggregate GPU time than the most accurate single-predictor setting in the compute benchmark and approximately 8.4-fold lower latency under parallel execution. The main source of this gain was broader candidate coverage: the oracle ceiling rose from 64.1% for 100 OpenDDE structures to 70.5% for the 5 × 3 pool, while the respective ranking procedures recovered similar fractions of those ceilings. Diversity therefore creates the opportunity for improvement, and `AbPoolRank` is required to realize it. This conclusion was not specific to our scoring function: VoroIF-GNN and DeepRank-Ab also benefited from mixed pools, whereas DProQA, whose scores tracked DockQ poorly, did not. On identical pools, `AbPoolRank` achieved the highest raw Success@1, matched the strongest learned rankers within statistical uncertainty, outperformed the consensus methods and required only about one second per pool on a CPU.

Several results clarify what makes diversification useful. Predictor value was governed by complementarity rather than standalone accuracy: Boltz-2 had a relatively low single-predictor oracle ceiling but contributed strongly to high-performing subsets, whereas a three-predictor pool of Boltz-2, Promera and OpenDDE matched the full pool at substantially lower compute. `AbPoolRank` also transferred across predictor distributions without model-specific calibration. Leaving any one development predictor out of training reduced performance by at most 3 percentage points, and predictors unseen during fitting could still improve the pool at inference. More stringently, full MSA-conditioned ESMFold2 was introduced only after method development; the frozen `AbPoolRank` model accepted it without retraining or loss of accuracy. Adding this external predictor did not further improve Success@1, however, despite a small increase in oracle coverage. Together with `AbPoolRank`’s tendency to favour the highly self-consistent OpenDDE distribution (Supplementary Note 2), this exposes an important limitation of consensus-based ranking: agreement is an imperfect proxy for correctness when an accurate predictor samples structurally unusual poses. Separating cross-source agreement from within-source self-consistency may therefore provide a route to better selection, particularly as pools become larger and more heterogeneous.

Several limitations define the scope of these conclusions and point to the next experiments. First, the study is restricted to antibody–antigen complexes, where predictor confidence remains imperfect and where different co-folders often explore distinct binding modes; whether the same breadth-over-depth advantage holds for general protein–protein complexes remains to be established. Second, although `AbPoolRank` matched or exceeded the strongest external rankers, substantial oracle headroom remained: high-quality structures were present in 48.0% of mixed pools but selected in only about 31% of cases. Improving recognition of high-quality but weakly supported poses is therefore a major opportunity, particularly for predictors that are accurate yet structurally idiosyncratic. Finally, the relative value of individual predictors depended on the composition and sampling depth of the pool, suggesting that future systems should adaptively allocate compute across complementary sources rather than treat all predictors as equally informative. Broader evaluation across complex classes, larger predictor panels and adaptive source selection will determine how generally this strategy works.

## Acknowledgments

This work was supported by the National Institutes of Health under award number 1R35GM141861 and Sanofi, including a research grant from Large Molecule Research as a 2025-26 Open Innovation Lab award.

## Data availability

Antibody-antigen complexes and cluster splits were obtained from SAbDab2 (https://zenodo.org/records/20083995) [27].

## Code availability

Source code is available at https://github.com/VarunUllanat/abpoolrank.

## Authors contributions

V.U., A.G. and R.L. conceptualized the project. V.U. generated and processed the data, ran training, and carried out downstream evaluation. P.A., N.F. and B.B. led the study. All authors contributed to writing the manuscript.

## Methods

### Task formulation

Given an antibody–antigen target *t* and a pool of candidate co-folded structures *P*_*t*_ = {*x*_1_, …, *x*_*N*_ *}*drawn from one or more predictors, a selector produces scores *s*_*i*_ and returns 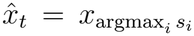. True labels *q*_*i*_ = DockQ(*x*_*i*_) [28] are never observed at inference. The deployment question is not which predictor is best, but how a fixed budget of *B* structures should be distributed across predictors and how one should be chosen from the resulting pool.

### Dataset and partition

Structures derive from the SAbDab2 machine-learning release [27]. The partition is a temporal, antigen-aware cluster split that uses the released antibody–antigen connected components as the indivisible unit, so no cluster spans the split; cluster date is the minimum deposition date over all cluster members. Only protein antigens are retained (peptide antigens are excluded because DockQ is noisier on very short antigens); multi-chain antigens are retained and every antigen chain contacting the antibody is preserved. Both paired antibodies and single-domain VHH antibodies are included, and each target’s pool is homogeneous. The delivered cohorts comprise 614 training complexes and 106 test complexes.

Four predictors contributed exactly 100 labeled poses for all 106 test targets and one contributed 100 poses for 103; all analyses therefore use the 103-target complete-case cohort in which every predictor contributes exactly 100 poses. Every method, baseline and oracle within a comparison selects from identical candidate identifiers, so no arm is evaluated on an easier subset than another.

### Structure generation

Five co-folding predictors were run, each in an isolated environment behind a thin wrapper consuming a common input specification: Boltz-2 (*boltz* 2.2.0) [23], Protenix-v2 (*protenix* 2.0.0) [24], Promera [5], ESMFold2-Fast (*esm* 3.3.0, *biohub/ESMFold2-Fast*) [25] and OpenDDE (*opendde* 1.0.2) [26]. A sixth predictor, ESMFold2 (*biohub/ESMFold2*, 20 trunk loops, MSA encoder enabled), was generated on the test targets only and is used exclusively as a fully external evaluation predictor.

Every predictor used the same protocol: *S* independent seeds ×5 diffusion samples per seed, with *S* = 20 on test (100 poses per predictor per target) and *S* = 4 on training (20 poses). Seeds were frozen before generation. Structural template retrieval was disabled in every wrapper. Multiple sequence alignments are per-chain a3m files from a single internal ColabFold [29] MMseqs2 server [30], cached so that all MSA-conditioned predictors receive identical alignments. ESMFold2-Fast is single-sequence and receives none.

Predicted chains are relabeled H (heavy or VHH), L (light, when present) and by manifest identifier for antigen chains, mapped by sequence alignment rather than by predictor-specific chain labels. The antibody is {H, L} and the antigen is every remaining chain; H–L and antigen–antigen contacts are excluded from all interface features and labels.

### Quality labels

DockQ 2.1.3 was used as ground-truth labels. It is is computed over all native interfaces, restricted to interfaces whose two chains cross the antibody–antigen boundary, and averaged over those interfaces. DockQ’s global summary over all interfaces is deliberately not used as the label, because it folds in antigen–antigen and heavy–light contacts and inflates the score. When no antibody–antigen interface is found the label is null rather than being silently replaced.

### Metrics and statistics

The primary metric is Success@1, the fraction of targets whose selected structure attains DockQ ≥0.49 (CAPRI medium or better), reported in percentage points. Success@1 was chosen over mean DockQ because mean DockQ rewards improvements that cross no decision boundary.

At total budget *B* over *G* predictors, each contributes ⌊*B/G*⌋ poses with the remainder spread over the first predictors. Each (target, budget) pool is redrawn 100 times and the pertarget Success@1 is the mean over those draws; pose sub-sampling contributes at most 0.15% of total variance at 100 repeats, so the reported uncertainty is dominated by the 103-target cohort rather than by the draw.

The resampling unit is always the biological target as pose-level resampling is never used. All method comparisons are paired by target: per-target Success@1 is computed for both arms on the intersection of their target sets and the per-target difference is bootstrapped (20,000 resamples for the comparisons in Figs. 1 and 2, 4,000–10,000 for the transfer, composition and control analyses).

### Pose-pair descriptors

For every target, one symmetric *N* ×*N* matrix per channel is precomputed over all candidates. No native structure is read. A descriptor is computed once per pose from heavy atoms—the epitope set (antigen residues within 5 Å of the antibody), the paratope set, the antibody–antigen contact-residue-pair set, and a soft epitope weighting *w* = *σ*((5 −*d*)*/*1)—and then all pairs are formed after superposing the two poses on their shared antibody C*α* atoms by the Kabsch algorithm, requiring at least ten shared residues. Seven channels are stored: epitope, contact-pair, paratope and soft-epitope Jaccard indices; geom sim = 1*/*(1 + RMSD) of antigen C*α* atoms after antibody alignment; antigen center-of-mass displacement; and relative antigen rotation angle. These are all prediction-to-prediction quantities that use predictions as pseudo-references.

### AbPoolRank design

All consensus statistics enter as *within-pool percentiles*, pct(*v*)_*i*_ = rank(*v*_*i*_)*/*(*n*− 1). This is necessary because absolute consensus similarity is systematically higher on training than test targets (median geom sim 0.134 versus 0.036), as training complexes lie inside the predictors’ training windows and their pose distributions collapse. Absolute similarity therefore does not transfer but its within-pool rank does.

Given a similarity matrix and source labels *g*_*i*_ over *G* sources, pose-similarity support is 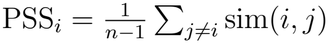sim(*i, j*) and source-balanced support caps each other predictor at one vote, 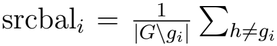, 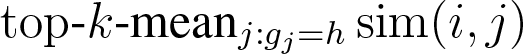 with *k* = min(3, *m*). `AbPoolRank` reads four features: the within-pool percentiles of PSS and of source-balanced support on geom sim, the percentile of PSS on epitope Jaccard, and log *N*. Widening the model to all seven pose-pair channels did not change performance measurably (Supplementary Note 3).

Exactly one feature uses source information: source-balanced support requires knowing which poses share a source—the partition, never which model produced it. The accurate description is therefore pose geometry plus an anonymous source partition, and a similarity-only ablation that removes even that quantifies its contribution. The number of sources cannot reach the features through this normalization: the factor 1*/*| *G*\ *g*_*i*_ |is the same constant for every pose in a pool, so it cancels exactly from the percentile, which we confirmed at *G* = 4, 5 and 6. This is why a selector trained at one pool composition transfers to another.

### Training

The regression target is the within-pool percentile of DockQ, not DockQ itself. Fitting absolute DockQ is misspecified here since training pools average 0.467 DockQ against 0.283 on test (since predictors are trained on these pre-2021 complexes). Training pools are formed by drawing *m* ∈ { 1, 2, 4, 10, 20} poses per predictor. `AbPoolRank` uses two independent draws per (target, *m*) over the 608 training complexes for which every predictor supplied structures (5,368 pools); `AbPoolRank` refitted inside the transfer, composition and control analyses use three draws per (target, *m*) (8,052 pools), so that all arms within each of those analyses share one training procedure.

The model is a histogram gradient-boosted regressor (250 iterations, learning rate 0.07, minimum 40 samples per leaf). Three models are fitted with different random seeds and their predictions averaged, because the threaded histogram build is not reproducible.

### Baselines

Native confidence uses each predictor’s own documented ranking quantity, verified per predictor rather than assumed: Boltz-2 reports a composite confidence score [23], Promera reports an interface-contact probability [5], and Protenix-v2, ESMFold2-Fast, OpenDDE and ESMFold2 report ipTM. Single-predictor baselines take the argmax of that quantity over the predictor’s own structures to get Success@1, which is the standard deployment procedure. On a mixed pool the same argmax is reported as a floor. The oracle is the best structure present in the pool and is not a deployable method.

### External rankers

Seven external rankers (six published methods, plus a DockQ-based jury) were run on the same structures without retraining or parameter changes. Each received the complete structures with the antibody as chains H and L (H only for VHHs) and the antigen as the remaining chains; each method’s top-ranked structure was credited with its DockQ. Single-structure rankers scored all 51,500 test structures once: VoroIF-GNN-v2 [20] (default model set, contacts restricted to antibody–antigen chain pairs, ranked by its pseudo-goodness score), DeepRank-Ab [11] (predicted DockQ; ESM-2 650M embeddings) and DProQA [19] (predicted DockQ). In DProQA, a structure whose secondary-structure assignment failed was skipped instead of aborting the whole target. Consensus methods were computed within each pool. Pcons [12] used its S-score, which by design does not compare structures from the same predictor. 3D-Jury [13] summed MaxSub similarities (number of C*α* pairs within 3.5 Å after superposition, MaxCluster 0.6.6; values below 40 set to zero) over the other pool members and divided by pool size plus one; a variant took, for each other predictor, only its most similar structure. MULTICOM PSS [15] averaged min(TM_1_, TM_2_) from US-align multimer alignment (version 20241108, `-mm 1 -ter 1`). The DockQ-jury averaged DockQ of a structure scored against each other pool member as reference, over antibody–antigen interfaces. Structures a method could not score (at most 10 of 51,500 for any single-structure ranker, structures without antibody–antigen contact or with a failed secondary-structure assignment) were ranked last. Runtime was measured on 20 randomly chosen 15-structure pools, one CPU core per method, with `AbPoolRank` model loaded once.

### Diversity controls

Two controls test whether the gain requires genuinely different predictors rather than the mechanics of `AbPoolRank`. In the *context-only* control, the pool contains structures from several predictors but the selection is restricted to those from one primary predictor, so any gain reflects cross-predictor agreement and cannot come from a better candidate entering the selectable set. In the *pseudo-source* control, 15 poses are drawn from a single predictor and split into five arbitrary groups of three, which are then presented to frozen `AbPoolRank` as five sources; total pool size, number of sources and poses per source match the five-predictor pool exactly, and only the diversity is artificial. Group assignment is a seeded random partition, and the control is evaluated over the same 100 pool draws per target as the corresponding real pools, so all comparisons are paired within a run.

### Transfer experiments

For each held-out predictor, a `AbPoolRank` version is trained on training pools containing only the other predictors, frozen, and applied to test pools containing the held-out one; it is never refitted and never receives that predictor’s confidence, identity or labels. Two distinct leave-one-out experiments are reported and must not be conflated: removing a predictor from the selector’s *training data* while testing on pools that contain it (a transfer question), and removing a predictor from the *inference pool* (a deployment question).

Whether frozen `AbPoolRank` exploits an unseen predictor is assessed at fixed total budget, so that adding a source necessarily reduces sampling depth on the others; the oracle is reported for both pools, because a selector that ignored an unfamiliar source would leave realized success flat while the ceiling rose. The fully external predictor is the strictest form of this test, since it contributed no training data of any kind.

### Pool composition

All 31 non-empty predictor subsets were evaluated at a fixed 15-structure budget over 100 pool draws per target, with the oracle. Subset choice uses split-half nested selection: the best subset of each size is chosen on a random half of the 103 targets and scored on the other half, over 4,000 splits.

### Compute measurement

Wall-clock time was measured on one exclusive NVIDIA A100 80 GB GPU with cached MSAs. A cost model *C*_*g*_(*N*) = *A*_*g*_ + *s*_*g*_ ·seeds + *d*_*g*_ ·*N* isolates fixed setup, per-seed and per-sample cost, fitted only on targets measured in all configurations. A pool pays *A*_*g*_ once per included predictor. Aggregate GPU-hours and wall-clock latency under parallel execution are reported separately, since the latter uses more hardware rather than less compute.

