## Supplemental text for "Breadth over Depth: Predictor-Agnostic Consensus for Improved Antibody–Antigen Structure Prediction"

This PDF file includes:

Supplementary Notes 1 to 3

Supplementary Figures S1 to S5

Supplementary Table S1

### Supplementary Note 1: pool composition reflects complementarity rather than standalone accuracy

We compared all 31 non-empty predictor subsets at a fixed 15-structure budget to determine which combinations retained the benefit of pooling (Supplementary Fig. S5a). To evaluate subset choice across targets, we performed 4,000 random split-half replicates: for each subset size, we selected the best predictor combination on one half and evaluated it on the other, recording how frequently each predictor was included. Subset selectors were refitted with a common procedure and evaluated on 100 pool draws per target; the full five-predictor pool therefore scores 58.2% here, against 58.0% for the released selector in Fig. 1.

Averaged over all subsets of a given size, Success@1 increased monotonically from 33.0% with one predictor to 58.2% with all five, while oracle coverage increased from 43.7% to 70.0%.

The parallel increases in realized and oracle performance support a coverage benefit from adding predictors. However, performance varied substantially among subsets of the same size. The average benefit of breadth therefore coexisted with specific smaller combinations that were especially effective; selecting such combinations requires information about complementarity, not only individual accuracy.

Boltz-2 was the clearest example. Its single-predictor oracle ceiling at the matched budget, 32.9%, was among the lowest (ESMFold2-Fast, 32.4%), yet it appeared in 71% of selected two-predictor subsets and 99% of selected three- and four-predictor subsets (Supplementary Fig. S5b). Protenix-v2 had much higher standalone oracle coverage, 53.4%, but appeared in fewer than 1% of selected pairs. A low individual oracle ceiling describes how often a predictor supplies a successful candidate; it does not describe how much those successes overlap with the candidates supplied by other models. Boltz-2's inclusion pattern was consistent with the pairwise complementarity analysis, in which every significant pairwise gain involved Boltz-2 or Promera (Supplementary Fig. S4a). Its value in a mixed pool was therefore substantially greater than its standalone ranking would suggest.

OpenDDE represented a different contribution. It appeared in at least 99.9% of selected subsets of two to four predictors, consistent with its strong standalone performance, despite limited pairwise complementarity with Protenix-v2 and ESMFold2-Fast. Protenix-v2 and ESMFold2-Fast were selected least frequently. These membership patterns reflect performance-based choice of predictor combinations; the structural selector itself still receives only anonymous source groups. The results distinguish two reasons to retain a co-folder: consistently strong candidates, as with OpenDDE, and complementary predictions that add value beyond a model's individual coverage, as with Boltz-2.

We also removed each predictor from the full inference pool while holding the trained selector fixed and redistributing the 15-structure budget across the remaining sources (Supplementary Fig. S5c). Unlike exclusion during fitting, this ablation changes the candidates and structural context available at deployment. Removing OpenDDE reduced Success@1 by 9.6 pp (95% CI, 4.7 to 14.8), followed by Boltz-2 at 7.3 pp (3.0 to 12.1) and Promera at 2.1 pp (0.4 to 4.2); the effects of removing Protenix-v2 or ESMFold2-Fast were smaller and not resolved. The particularly large Boltz-2 effect reinforces the distinction between individual accuracy and marginal pool value. Because a source contributes both selectable structures and support for structures from other sources, removal can affect both coverage and consensus. Pool membership should consequently be evaluated through the performance of the combined system, rather than inferred directly from a standalone leaderboard.

### Supplementary Note 2: which predictor supplies the selected structure

The marginal value of adding a predictor declined as the sampling budget grew, both for the five leave-one-out predictors and for the fully external one. To characterise this we recorded, for pools of four structures from each of the six predictors, which source supplied the selected structure and which supplied the oracle-best structure, over 30 draws on each of the 103 targets.

| Source | Held oracle-best (%) | Selected (%) | Mean support percentile |
|---|---|---|---|
| OpenDDE | 25.7 | 45.5 | 0.677 |
| Boltz-2 | 20.2 | 17.9 | 0.461 |
| Protenix-v2 | 19.1 | 15.1 | 0.548 |
| ESMFold2 | 14.8 | 8.9 | 0.488 |
| Promera | 11.7 | 7.0 | 0.427 |
| ESMFold2-Fast | 8.6 | 5.5 | 0.398 |

The selector chose OpenDDE structures roughly twice as often as OpenDDE held the oracle-best candidate, and chose every other source less often than that source held the best candidate. The final column offers a plausible account: OpenDDE attains by far the highest mean within-pool percentile of pose support, and its own structures are the most mutually similar of any predictor (within-source similarity 0.2588, against 0.1413 for ESMFold2 and 0.1062 for ESMFold2-Fast). A selection rule based on agreement will favour the source whose structures agree most, which need not be the source that is most often correct. Deeper pools give such a source more opportunities to dominate the ranking, which is consistent with the declining marginal value of additional predictors, although these observational data do not establish that mechanism. A support statistic that separated agreement with other sources from internal self-consistency, rather than combining them, is a natural direction for future work.

### Supplementary Note 3: feature-set ablation

The selector reads two of the seven cached pose-pair channels. Widening it changes nothing measurable. Fitting the same model on identical pools with pose-similarity support over all seven channels (eight features), or with both support statistics over all seven (fifteen features), gave paired differences against the four-feature model of $+0.2$ to $+1.1$ and $-0.1$ to $+1.5$ percentage points respectively, with every interval spanning zero at every budget. The additional channels are used—permutation importance ranks the source-balanced rotation term second, behind geometric pose support—but they are redundant: antigen centre-of-mass displacement and relative rotation are components of the same rigid-body transform that the geometric similarity channel already summarises, and the four Jaccard indices are alternative views of one contact-set overlap. Two channels, one geometric and one epitope-based, span the available signal, and the four-feature model is retained on grounds of simplicity.

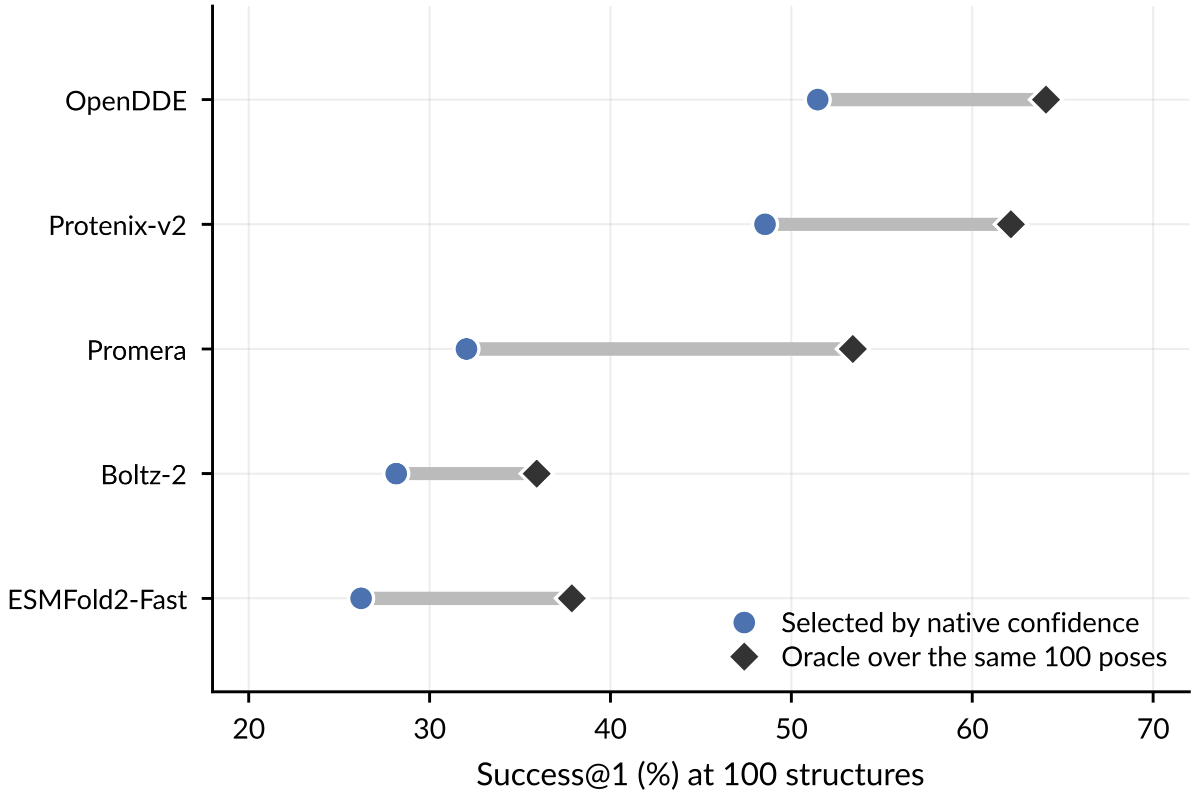


**Figure S1: Native confidence leaves substantial headroom within every predictor.** Per-predictor Success@1 of native-confidence selection against the oracle ceiling of the same 100-structure pool, on the 103-target cohort. The gap is quality present in the pool but not identified by the predictor's own confidence, and motivates treating selection rather than generation as the binding constraint.

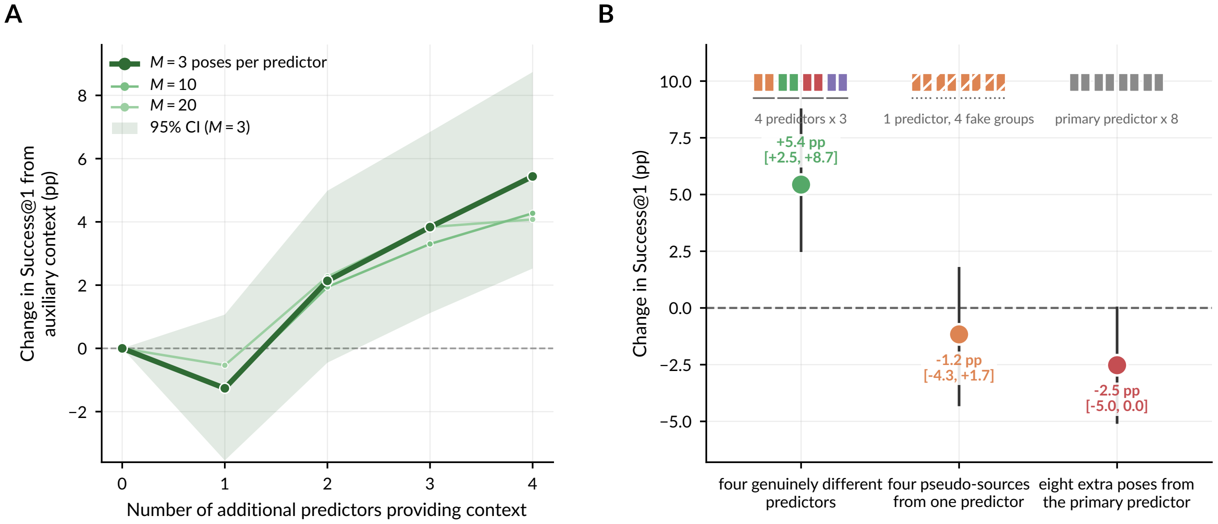


**Figure S2: Context-only control.** Success@1 when a fixed number of candidate structures from one primary predictor is ranked with additional structures supplied only as context, so that the selected pose always originates from the primary predictor. This isolates the contribution of cross-predictor *agreement* from the contribution of cross-predictor *coverage*: any gain here cannot come from a better candidate entering the pool.

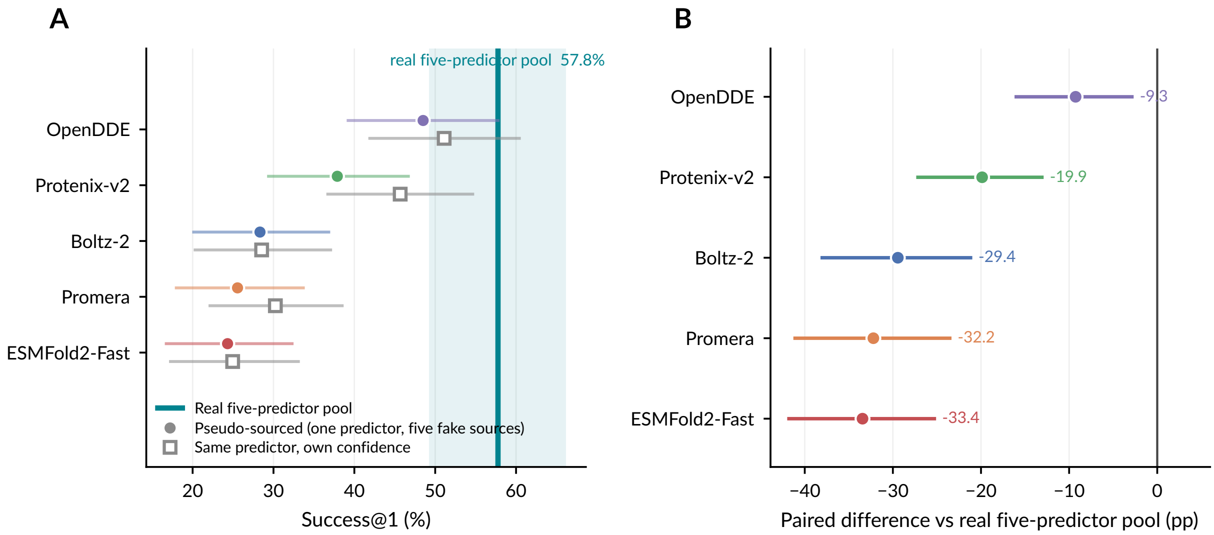


**Figure S3: Pseudo-source control: artificial source partitioning does not reproduce the gain.** Fifteen poses are drawn from a *single* predictor and split into five arbitrary groups of three, which are then presented to the frozen selector as five sources. Total pool size, number of sources and poses per source match the real five-predictor pool exactly; only the diversity is artificial. The real pool is scored here by the selector refitted for this control on the control's own pool draws (57.8%), so that every comparison is paired; the released selector reaches 58.0% on the draws of Fig. 1. **a,** Success@1 of each pseudo-sourced pool (filled circles) against the same predictor's own confidence over the same 15 poses (open squares), with the real five-predictor pool as the vertical reference and its marginal interval shaded. Fake partitioning neither approaches the real pool nor improves on the predictor's own confidence. **b,** Paired difference between each pseudo-sourced pool and the real five-predictor pool on the same targets; bars are 95% paired-bootstrap intervals. The source-balanced feature therefore does not manufacture an advantage from partition structure alone.

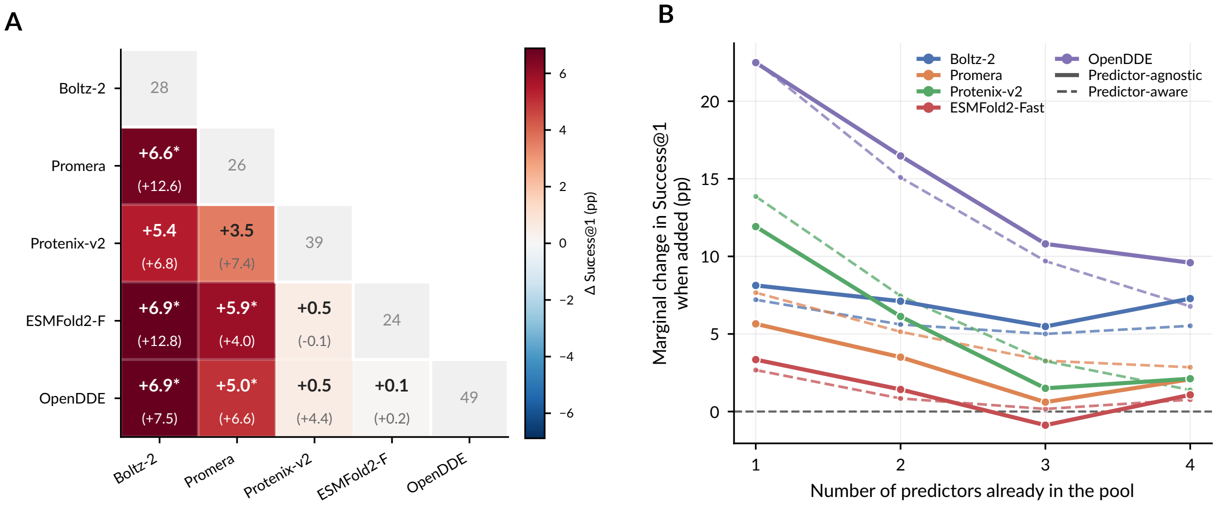


**Figure S4: Pairwise complementarity and marginal value, under the predictor-agnostic selector. a,** For each predictor pair at a 15-structure budget, Success@1 of the pair minus that of its stronger member; realized values in bold with the oracle in parentheses, and the diagonal giving each predictor alone. The gap between realized and oracle is coverage the pair contains but selection misses. **b,** Marginal change in Success@1 when each predictor is added to a pool of a given size, at a fixed 15-structure budget. Both panels are computed under the deployed predictor-agnostic selector, so they describe complementarity as the primary method actually realizes it.

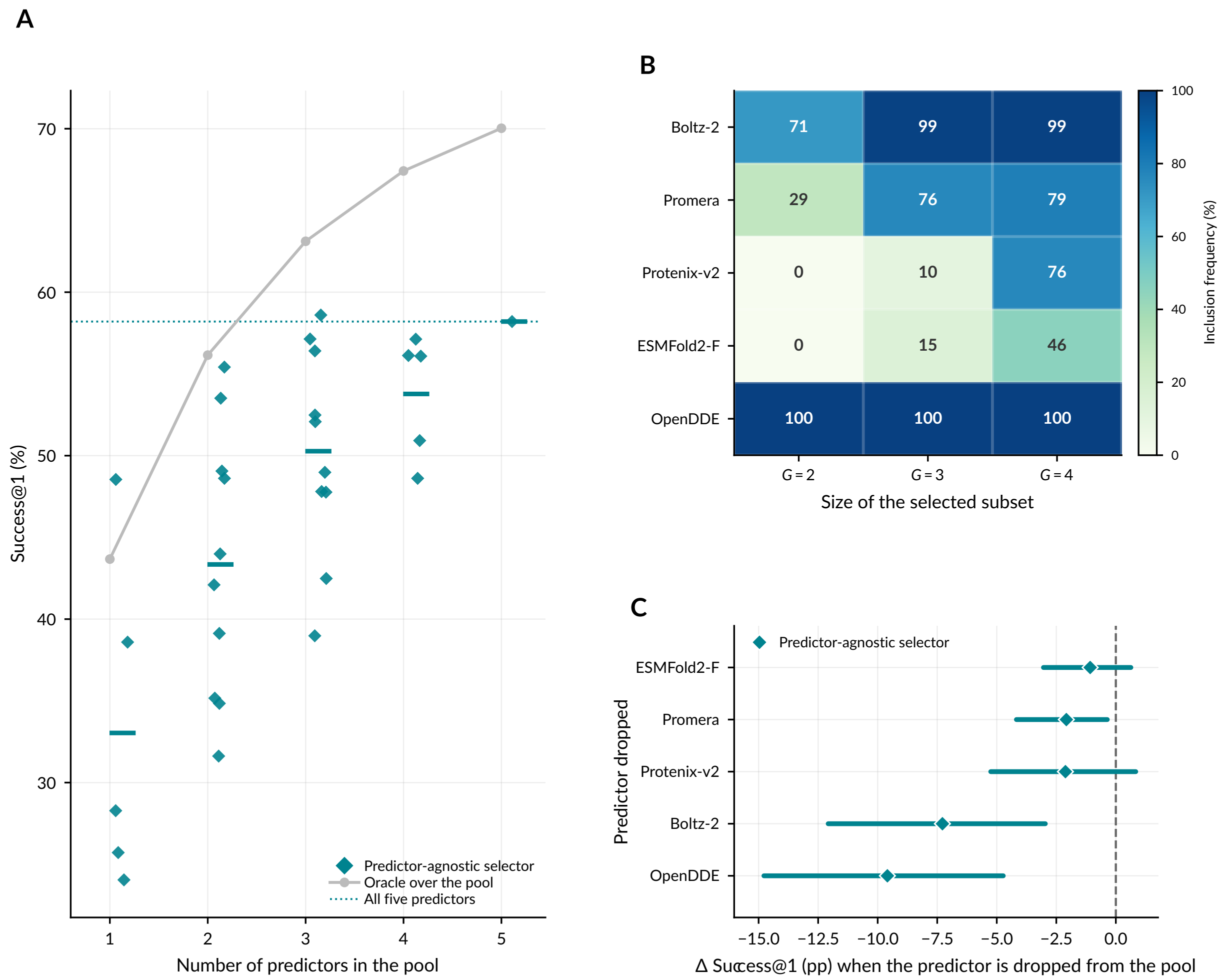


**Figure S5: Pool composition reflects complementarity rather than standalone accuracy. a,** All 31 non-empty predictor subsets at a fixed 15-structure budget. Each point is one subset; horizontal bars are means within a pool size. Teal diamonds, the predictor-agnostic selector; grey line, the pool oracle; dotted line, the full five-predictor pool. **b,** Inclusion frequency under split-half nested selection (4,000 replicates): the best subset of each size is chosen on one random half of the 103 targets and its composition recorded before evaluation on the other half. Frequencies are for the predictor-agnostic selector, which is never given predictor identities, so the composition shown emerges from pose geometry alone. **c,** Paired change in Success@1 when one predictor is removed from the inference pool at 15 structures. This is the deployment question—which predictors must a user actually run—and is distinct from Fig. 3b, where a predictor is removed from the selector's *training* data.

**Table S1: The predictor panel.** Versions, declared structural training cutoffs and native confidence fields for the six co-folding predictors, with per-predictor performance on the 103-target complete-case cohort. Mean DockQ is over all 100 sampled structures per target. Top-1 is Success@1 using that predictor's own native confidence over its 100 structures; the oracle is the best structure available in the same 100. Full MSA-conditioned ESMFold2 was generated on the test targets only and is used solely as a fully external evaluation predictor, so it contributes no training data of any kind.

| Predictor | Version | Cutoff | Confidence field | Mean DockQ | Top-1 (%) | Oracle (%) |
|---|---|---|---|---|---|---|
| OpenDDE | 1.0.2 | 2021-09-30 | ipTM | 0.397 | 51.5 | 64.1 |
| Protenix-v2 | 2.0.0 | 2021-09-30 | ipTM | 0.305 | 48.5 | 62.1 |
| ESMFold2* | esm 3.3.0 | 2021-09-30 | ipTM | 0.270 | — | — |
| Boltz-2 | 2.2.0 | 2021-09-30 | confidence score | 0.265 | 28.2 | 35.9 |
| Promera | source | 2023-12-31 | interface-contact prob. | 0.223 | 32.0 | 53.4 |
| ESMFold2-Fast | esm 3.3.0 | 2021-09-30 | ipTM | 0.203 | 26.2 | 37.9 |

*Full MSA-conditioned ESMFold2 (`biohub/ESMFold2`, 20 trunk loops, MSA encoder enabled), as distinct from ESMFold2-Fast (`biohub/ESMFold2-Fast`, MSA encoder disabled, 3 trunk loops). The two are separate checkpoints and are never pooled under a single source label; their mean pairwise pose similarity (0.0801) ranks sixth of fifteen cross-predictor pairs, below Protenix-v2 with OpenDDE (0.1342). Top-1 and oracle entries are omitted because this predictor is evaluated only as an addition to pools, not as a standalone arm.
